# The Respiratory-Pupillary Phase Effect: Pupils size is smallest around inhalation onset and largest during exhalation

**DOI:** 10.1101/2024.06.27.599713

**Authors:** Martin Schaefer, Sebastiaan Mathôt, Mikael Lundqvist, Johan N. Lundström, Artin Arshamian

## Abstract

The prevailing view in the study of animal and human pupillary function has been that pupils dilate and are largest during inhalation and constrict and are smallest during exhalation. However, this notion has recently been challenged. Here, we systematically address this question by conducting a study encompassing three experiments (two resting tasks and one visual perception task), with the latter two being pre-registered. Collectively, across nasal and oral breathing, resting, and task conditions, our experiments consistently and robustly demonstrate that pupil size is smallest around inhalation onset and largest around peak exhalation. This phenomenon, which we term the Respiratory-Pupillary Phase Effect (RPPE), directly contradicts the long-held notion that pupils are largest during inhalation and smallest during exhalation. Notably, the dilation and constriction processes overlap with both phases. The observed consistency and significance of the RPPE across various conditions underscore the need for further investigation into its underlying mechanisms and potential impact on human behavior.

## Introduction

For over half a century, the prevailing view in the study of animal and human pupillary function has been that pupils dilate and are largest during inhalation and constrict and are smallest during exhalation (Ashhad et al., 2022; Borgdorff, 1975; Golenhofen & Petranyi, 1967; Loewenfeld, 1958). This claim has been widely accepted as a basic component of mammalian physiology (Schaefer et al., 2022). However, a recent systematic review on human respiration and pupillary function by Schaefer et al. (2022) challenged this long-held notion. Specifically, the empirical support for these claims was shown to be scarce and at best, inconclusive with conflicting findings (e.g., Borgdorff, 1975; Nakamura et al., 2019; Schaefer et al., 2022). This lack of evidence primarily stems from methodological limitations including small sample sizes, inadequate or absent statistical analysis, inadequate physiological measurements, and failure to control for eye blinks (Schaefer et al., 2022). Here we address these limitations by re-investigating the link between breathing cycle and pupil size. We report strong evidence that—in contrast with the prevailing view—pupils are smallest around inhalation onset and largest during exhalation with the dilation and constriction processes overlapping both phases.

Notably, the primary experimental evidence supporting the notion that pupils are largest during inhalation does not involve humans; instead, it originates from a single study conducted on cats (Borgdorff, 1975). This foundational study claimed that pupil dynamics are synchronized with the respiratory rhythm, showing maximum dilation during inhalation and minimal dilation during exhalation. However, this phenomenon was only observed in cats that were lightly anesthetized or tranquilized, with the effect completely disappearing in both fully awake as well as heavily anesthetized cats. While Borgdorff (1975) clearly stated the limitation of this local phenomenon, it has since been misrepresented in the scientific literature as demonstrating a general physiological mechanism by which the two respiratory phases (inhalation and exhalation) shape pupil dynamics. This misinterpretation is particularly unfortunate considering the growing body of evidence highlighting the potential significance of both respiration and pupil function, and their potential interaction, in brain and behavior research (Arshamian et al., 2018; Heck et al., 2017; Ito et al., 2014; Joshi & Gold, 2020; Kluger et al., 2021a; Mathôt, 2018; Perl et al., 2019; Schwalm & Rosales Jubal, 2017; Tort et al., 2018; Zelano et al., 2016). For example, breathing creates neural oscillations that propagate globally (e.g., visual cortex and hippocampus), directly influencing behavior (Heck et al., 2017; Herrero et al., 2017; Kluger et al., 2021a; Tort et al., 2018). During both nasal and oral inhalation these oscillations are mainly created by the brainstem Pre- Bötzinger complex (Negro et al., 2018). However, nasal breathing also includes an additional oscillator—the olfactory bulb. Olfactory sensory neurons at the roof of the nasal cavity detect mechanical pressure caused by airflow in the nostrils, even in the absence of odors (Grosmaitre et al., 2007), and project this information to the olfactory bulb, which in turn generates brain oscillations that propagate globally and may uniquely affect perceptual and cognitive functions (Arshamian et al., 2018; Fontanini & Bower, 2006; Heck et al., 2017; Karalis & Sirota, 2022; Kay et al., 2009; Kocsis et al., 2018; Negro et al., 2018; Schaefer et al., 2024; Tort et al., 2018). This indicates that nasal and oral breathing might shape pupil dynamics differently. While breathing influences overall brain function and behavior, pupil size fluctuations provide a dynamic readout of brain and behavioral states (Mathôt, 2018; Schwalm & Rosales Jubal, 2017). Therefore, it is important to determine how respiration affects pupil dynamics within the fundamental tenets of mammalian physiology.

We systematically address this question by conducting a study encompassing three experiments including healthy participants, of which the latter two experiments are pre-registered. In Experiment 1, we first measured both respiratory patterns and pupil size in 50 subjects at rest under two breathing conditions: nasal and oral breathing. To the best of our knowledge this is the first-time pupil function has been studied as function of respiratory route (Schaefer et al., 2022). Studying participants at rest allowed us to determine the potential relationship between respiratory cycles and pupillary dynamics without the confounding influence of visual tasks or manipulations. To ensure the robustness of our initial exploratory findings, we pre-registered the analysis steps and results from Experiment 1 and then directly replicated them in an independent group of 53 new participants in Experiment 2. Finally, in Experiment 3 we determined whether the results from the two experiments at rest, generalized to active visual perception and assessed potential breathing-route-dependent differences. This involved testing 112 subjects who performed a visual perception task, once while breathing nasally and once orally.

Collectively, across nasal and oral breathing, resting, and task conditions, our experiments consistently and robustly demonstrate that pupil size is smallest around inhalation onset and largest around peak exhalation. This phenomenon, which we term the Respiratory-Pupillary Phase Effect (RPPE), directly contradicts the long-held notion that pupils are largest during inhalation and smallest during exhalation. Notably, the dilation and constriction processes overlap with both phases. The observed consistency and significance of the RPPE across various conditions underscore the need for further investigation into its underlying mechanisms and potential impact on human behavior.

## Materials & Methods

### Participants

We recruited a total of 112 participants (70 female) ranging in age from 18 to 42 years (mean age 28.0 years). Of these, 50 participants (35 female) ranging in age from 18 to 42 years, with a mean age of 27.9 years, were part of Experiment 1. Another 53 participants (30 female) ranging in age from 19 to 42 years with a mean age of 28.5 years, were part of Experiment 2. All 112 participants participated in Experiment 3. Three participants did not return for their second testing session (affecting Experiment 1 and 3).

All participants reported being healthy and able to breathe freely and independently through both their nose and mouth. Participants were instructed to refrain from ingesting caffeine on the day of the experiment as this is known to affect pupil size dynamics (Abokyi et al., 2017; Redondo et al., 2020). They received compensation for their participation in the form of movie ticket vouchers to the value of 360 SEK. Ethical approval was obtained from the national Swedish Ethical Review Authority (Dnr 2020-00972) and all participants signed informed consent prior to participation.

### Experimental setup

#### Experiment 1 and 2: At rest

Participants underwent two five-minute long sessions at rest, where they looked at a gray screen with a black fixation cross for five minutes. In one session, they only breathed through their nose, and in the other, only through their mouth. The order of which breathing route came first was alternated equally, and the sessions were conducted on separate days.

The participants were seated with their heads fixed in a chin rest in a dimly lit room, without any variation in light intensity through the experiments, at a distance of 61 cm from the screen. The participants were instructed to breathe normally and to keep their eyes open (blinking allowed).

#### Experiment 3: Visual task

We conducted a 3-alternative forced choice visual detection experiment (modelled after Kluger et al., 2021b) in the same luminance condition as during Experiments 1 and 2. During the task, participants fixated on a cross (0.4° in diameter) in the center of the screen presented against a gray background. Each trial consisted of a fixation period (jittered between 1200 and 3500 ms), followed by a brief (50 ms) presentation of a small (0.3° in diameter) Gabor patch in a marked circular area (3.5° in diameter) 10° to the left or the right side of the fixation cross, or no Gabor patch was presented (catch trials). After a delay of 500 ms, a question mark in the center of the screen prompted participants to press an arrow key to indicate whether they had seen a Gabor patch, and if so on which side of the screen. After the participants pressed one of the arrow keys, the next trial started. For each trial, target contrast was adapted by a QUEST staircase (Watson & Pelli, 1983) aimed at individual hit rates of about 60%. The task consisted of 762 trials with an equal number of left target, right target, and catch trials. The first 12 trials of the task were practice trials, and after every 30 trials the participants got a short ad libitum break. The behavioral results of this task are not part of this study and will be presented elsewhere. The pupil size recordings during the breaks were not analyzed as participants were free to remove their head from the chin rest and move around (while staying seated).

### Physiological measures

Breathing was measured in two ways. We used a non-restrictive breathing mask attached to a breathing cannula to obtain a direct measure of airflow (0202-1199, Tiga-Med, Germany). Additionally, we used a temperature probe (MLT415/D, ADInstruments, Colorado, US) attached to the inside of the breathing mask and recorded breathing by measuring changes in air temperature over time (ΔT℃). Using two breathing measures gave us a backup in case one of the measures malfunctioned and allowed us to use whichever measure contained less noise. The breathing route not assessed was covered topically with surgical tape to prevent participants from accidentally using the wrong breathing route. All breathing signals were amplified, digitized at 1000 Hz (LabChart 7.0, ADInstruments, Colorado, US), and low-pass filtered at 5 Hz (PowerLab 16/35, ADInstruments, Colorado, US). Pupil size was measured with a Gazepoint eyetracker with a sampling rate of 60 Hz and recorded with the Gazepoint software. Heart rate was measured in two ways to allow for additional validation of the data; using a finger pulse electrode, and a 3-leads ECG setup (LabChart 7.0, ADInstruments, Colorado, US).

### Preprocessing of data

#### Pre-registration

We conducted an exploratory analysis for Experiment 1. Based on those results, we preregistered the preprocessing and analysis steps for Experiment 2 on the Open Science Framework (https://osf.io/xwjqe). These preregistered steps were subsequently applied to Experiment 3 as well and are detailed below. The same analysis steps were then also used for Experiment 3, and are described in detail below.

#### Pupil data

The preprocessing of the pupil size data was done in several stages, separately for each eye, and mainly followed the suggestions of Mathôt and Vilotijević (2022).

First, any recording marked by the eyetracker as valid but shorter than 250ms in duration was marked as invalid to exclude data points with unreliable validity. In addition, valid data shorter than 500 ms which are surrounded by invalid data of more than 2 seconds, were also marked as invalid.

Second, any recording marked as invalid by the eyetracker for more than 500ms was removed to exclude invalid data that is unlikely to be blinks.

Third, any recording marked as invalid by the eyetracker or our above criteria was interpolated using the *fillmissing.m* Matlab function employing a piecewise cubic spline interpolation method. For each invalid data stretch, a buffer of two frames before and after was marked as invalid as well, and then the complete stretch was interpolated using the three valid samples preceding and following the buffer frames. If the range of the interpolated data points exceeded the range of the valid data points used for the interpolation by more than 30% (which might indicate that spikes have been introduced by the interpolation), the interpolation was redone using three, and then four buffer frames around the stretch of invalid data points. If the range of the interpolated data points still exceeded the range of the valid data points used for the interpolation by more than 30%, a linear interpolation using the 2 buffer frames and three valid samples around the stretch of invalid data points was used instead.

Fourth, the data was checked for rapid changes in pupil size that could indicate blinks or invalid interpolations. The stretches of data detected with this method were then interpolated in the same manner as described above, however; the cubic spline interpolation was only deemed as valid if its range did not exceed the range of the valid samples used for the interpolation by 20%. After finishing the interpolations, any valid stretches of data shorter than 250ms in duration were marked as invalid to exclude data points with unreliable validity. In addition, valid data stretches shorter than 500 ms which were surrounded by invalid data stretches of more than 2 seconds, and valid data stretches shorter than 1000 ms which were surrounded by invalid data stretches of more than 5 seconds were also marked as invalid.

Additionally, data values less or larger than the mean pupil size ± 3x the standard deviation of the pupil size values were marked invalid.

All interpolated pupil data was visually inspected to make sure the data looked reasonable, and participants with more than 30% invalid pupil data for both eyes after interpolation were excluded from further analysis.

After rejecting recording sessions with too much invalid pupil data, we were left with 38 complete participants (41 nose breathing sessions and 44 mouth breathing sessions) for Experiment 1, and 44 complete participants (48 nose breathing sessions and 47 mouth breathing sessions) for Experiment 2. For the visual detection task, we had 98 complete participants (103 nose breathing sessions and 102 mouth breathing sessions).

In line with our preregistration, we also assessed changes in absolute pupil size over time. To get the absolute pupil size values, we repeated the pupil interpolation in the same manner as for the pupil size values in pixels. However, the pupil size recordings in absolute values (mm) were less reliable than the recordings of the pupil size in pixels and we were therefore left with fewer valid recordings. After rejecting recording sessions with too much invalid pupil data, we were left with 30 complete participants (35 nose breathing sessions and 39 mouth breathing sessions) for Experiment 1, and 41 complete participants (47 nose breathing sessions and 43 mouth breathing sessions) for Experiment 2. The results for the absolute pupil size values are reported in the supplementary materials.

To increase power and reliability, we used the average of the left and right eye for each analysis where we looked at pupil size. Furthermore, we normalized all relative pupil size data (pixels) by z-scoring with mean zero and standard deviation of one, at the participant level to enable direct comparisons across participants.

#### Breathing data

Given that the airflow data exhibited a higher temporal sensitivity than the thermopod data, we opted to utilize the airflow data to determine the breathing phase for each participant. To analyze the respiratory patterns of each participant, we employed the BreathMetrics toolbox for Matlab, as outlined by Noto et al. (2018). This toolbox allowed us to identify key points in the breathing cycle, specifically the inhalation and exhalation onsets and peaks. These points of interest were then used to create a continuous measure of breathing phase spanning the full breathing cycle (360 degrees). This was achieved through linear interpolation: from 0° to 90° for the breathing samples between inhalation onset and inhalation peak, 90° to 180° from inhalation peak to exhalation onset, 180° to 270° from exhalation onset to exhalation peak, and 270° to 360° from exhalation peak to the subsequent inhalation onset. For subsequent analyses, aimed at detecting variations in pupil size across the breathing cycle, we divided the breathing cycle into 18 equally spaced bins, each spanning 20°, allowing us to compare pupil size during specific parts of the breathing cycle across participants. In- and exhalations lasting shorter than 500 ms or longer than 6000 ms were considered invalid (on average 7.15% of breathing data was considered invalid in this manner). Finally, the breathing data was downsampled to 60 Hz to match the sampling frequency of the pupil data.

### Data analysis

To investigate whether pupil size varied across the breathing cycle in a statistically significant manner, we calculated a two-way (breathing phase x breathing route) analysis of variance (ANOVA) comparing the average normalized pupil size values for each breathing bin and each breathing route.

We employed permutation testing to assess systematic patterns of pupil size fluctuations during the breathing cycle. For the nose and mouth breathing scenarios, we had an average of 35,879 and 34,381 observations respectively within the 20-degree breathing bins when combining the data of all participants in Experiment 1, an average of 44,027 and 41,440 observations per breathing bin in Experiment 2, and an average of 884,721 and 847,209 observations per breathing bin in Experiment 3. We randomly selected pupil size observations across all breathing cycle phases, equal to the calculated average number of observations per 20-degree bin. This random selection process was repeated 10,000 times, and during each iteration, the mean of the randomly selected pupil size observations was calculated. The resulting means from these iterations were graphed as histograms and compared against the mean pupil size value associated with each of the 18 unique breathing bins, and used to calculate *p*-values equal to two-sided t- tests (not corrected for multiple comparisons given our permutation assessment).

Statistical analysis were made in JASP (JASP Team, 2024) and Matlab. Figures were created in Matlab and esthetically modified in Inkscape (Inkscape Project, 2018). To show how pupil size changes across the breathing cycle, we created polar plots, polar histograms, and calculated the circular mean direction of individuals’ pupil size across the breathing cycle using the native *polarhistogram.m* and *polarplot.m* native Matlab functions, as well as the *circ_mean.m* function of the Circular Statistics Toolbox for Matlab (Berens, 2009).

## Results

Given the similarity of results from Experiments 1 and 2, we present the results of their combined data analysis for improved statistical power (individual experiment results can be found in the Supplementary Materials).

### Pupil dynamics at rest (Experiment 1 and 2 combined)

How individual participant’s pupil size fluctuated over the over the course of the breathing cycle varied (see Figure 1A). However, the average mean vector direction for all participants showed that for both nose and mouth breathing, the vast majority of the participants had largest pupils during exhalation (see Figure 1A).

**Figure 1.**
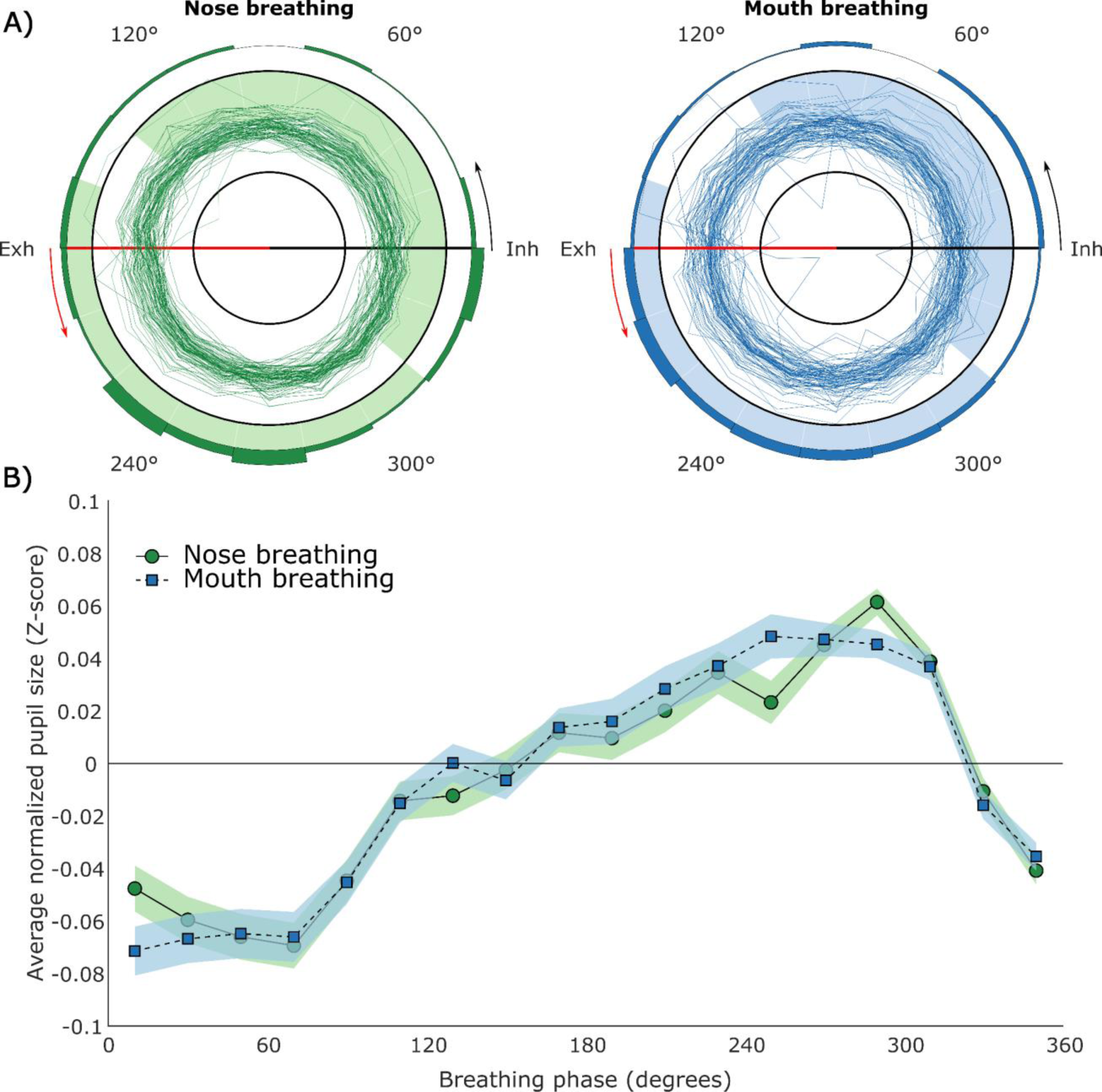
Pupil size at rest (5 minutes recording time). **A)** Polar plots of participants’ pupil size over the course of the breathing cycle. 0° (Inh) marks the onset of inhalation, and 180° (Exh) marks the onset of exhalation. The individual dark green/dark blue lines show the normalized pupil size values of individual subjects during nose and mouth breathing respectively. The inner black circle indicates a pupil size Z-score of -0.5, and the outer black circle indicates the pupil size Z-score of 0.5. The light green/light blue shading indicates a phase bin that is significantly different from the permutation testing results acquired by randomly sampling pupil size values from across the breathing cycle. A shaded bin outside the outer black circle indicates larger pupils, and a shaded bin on the inside of the outer black circle indicates smaller pupils than what could be expected from the permutation testing results. The polar histogram (dark green/dark blue bins) on the outside of the polar plot shows the proportion of circular mean directions pointing towards a certain breathing phase bin (the histogram bins add up to a total of 1). **B)** The line plot depicts the average normalized pupil size for each phase bin. Zero degrees on the x-axis corresponds to inhalation onset, and 180° corresponds to exhalation onset. The y-axis shows pupil size in Z-scores. The horizontal zero line represents the average pupil size for each individual. Pupil size during nose breathing is depicted by the green circles connected by a solid line, and pupil size during mouth breathing is depicted by the blue squares connected by a dashed line. The shaded areas represent the 95% confidence intervals.

The combined average of all participants while breathing at rest clearly demonstrated that pupil size peaked during exhalation and had a minimum during inhalation, for both nose and mouth breathing (see Figure 1B).

Furthermore, we statistically confirmed a significant effect of breathing phase (F(17) = 10.31, *p* < .001, ƞ^2^ = 0.070), but no significant effect of breathing route (F(1) = 1.14, *p* = .289, ƞ^2^ < 0.001), nor a significant interaction effect (F(17) = 0.50, *p* = .956, ƞ^2^ = 0.002).

The permutation testing revealed that 17/18 and 16/18 phase bins were significantly different from the randomly acquired distribution for nose and mouth breathing respectively, with 320° – 120/140° showing significantly smaller pupils, and 160° - 320° showing significantly larger pupils (see Table S1 and Figure 1).

### Pupil dynamics during visual task

Over a longer recording duration, participants’ pupil size fluctuation showed less variation over the course of the breathing cycle, and a clear, consistent pattern emerged (see Figure 2A). This convergence became especially clear when assessing the average mean vector direction for all participants, which showed that for both nose and mouth breathing, the vast majority of the participants had largest pupils around exhalation onset (see Figure 2A).

**Figure 2.**
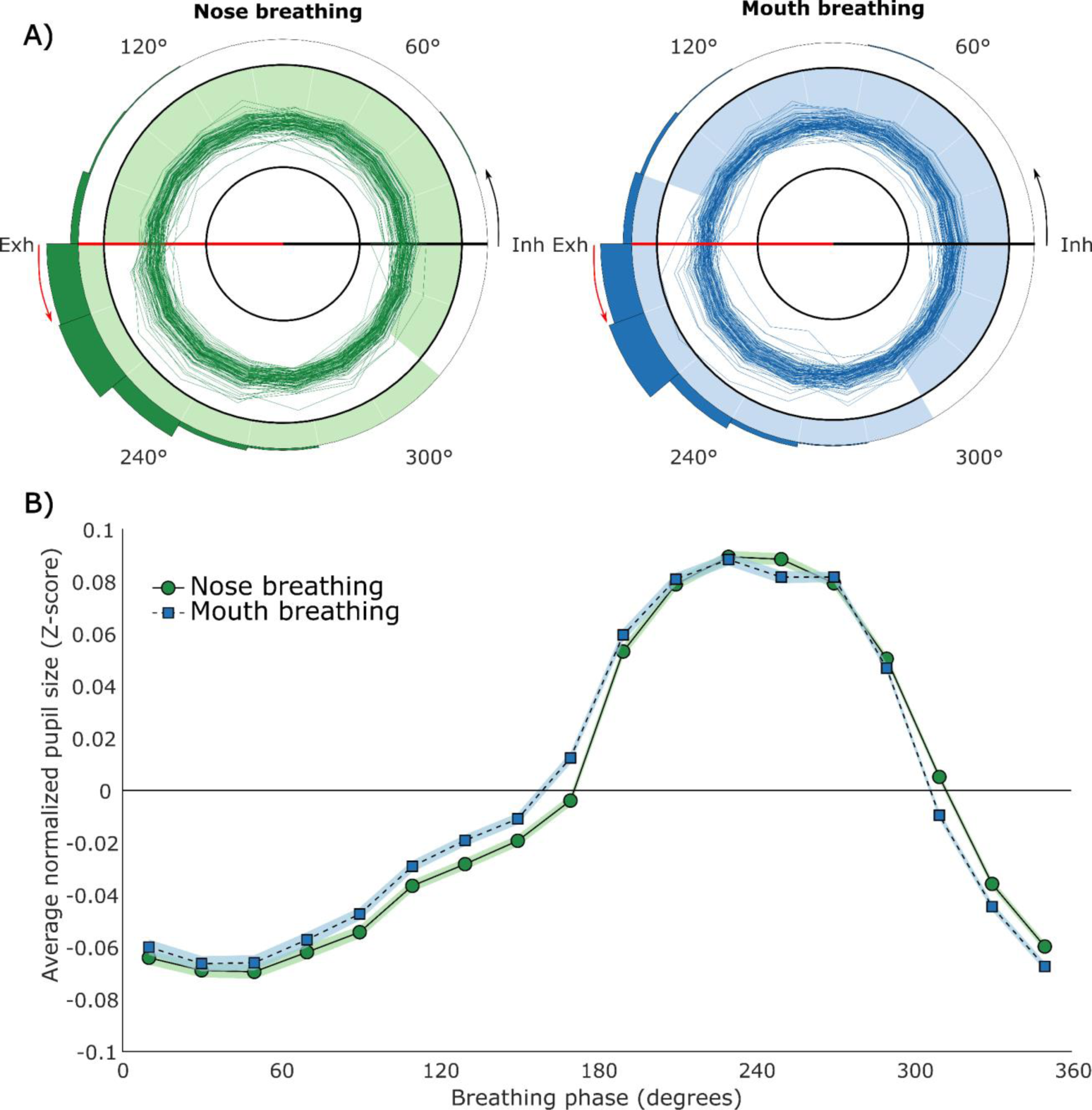
Pupil size during visual task (∼45 minutes recording time). **A)** Polar plots of participants’ pupil size over the course of the breathing cycle (see Figure 1A legend for a more detailed description). **B)** The line plot depicts the average normalized pupil size for each phase bin. Zero degrees on the x-axis corresponds to inhalation onset, and 180° corresponds to exhalation onset (see Figure 1B legend for a more detailed description). Finally, to highlight the difference between pupil size and changes in pupil size, we also plotted the change in pupil size over the course of the breathing cycle (see Figure 3). This plot reveals the ambiguous nature of previous claims that pupil size increases during inhalation and decreases during exhalation as in reality increases and decreases in pupil size occur during both breathing phases.

The combined average of all participants during the visual task again showed that pupil size tended to peak during exhalation and had a minimum around inhalation onset for both nose and mouth breathing (see Figure 2B).

Again, we statistically confirmed a significant effect of breathing phase (F(17) = 80.44, *p* < .001, ƞ^2^ = 0.365), but no significant effect of breathing route (F(1) = 3.43, *p* = .067, ƞ^2^ < 0.001), nor a significant interaction effect (F(17) = 1.00, *p* = .461, ƞ^2^ = 0.002).

The permutation testing revealed that all 18 phase bins were significantly different from the randomly acquired distribution for nose and mouth breathing, with 300/320° – 160/180° showing significantly smaller pupils, and 160/180° - 300/320° showing significantly larger pupils (see Table S1 and Figure 2).

## Discussion

In three experiments we demonstrate that pupil size systematically changes throughout the breathing cycle, across routes, and both at rest and during a visual task. Specifically, although dilation and constriction dynamics occur during both inhalation and exhalation, the pupil size consistently reaches its minimum around inhalation onset and its maximum during exhalation. This is in contrast to what was previously assumed based primarily on a single landmark study (Borgdorff, 1975). We term this phenomenon the Respiratory-Pupillary Phase Effect (RPPE).

Our study addressed limitations in prior research by employing a larger sample size, a preregistered replication, and by dividing the breathing cycle into 18 fine-grained bins, thereby enabling a more precise measurement of pupil size changes throughout the respiratory cycle compared to past binary inhalation-exhalation approaches (Schaefer et al., 2022). Notably, for the first time, we show that these changes in pupil size are not affected by breathing route, with nose and mouth breathing displaying the same pattern. The changes in pupil size we observed were highly significant yet relatively small. The modest size of this effect, combined with the small sample sizes used in prior research (median of 13 participants; Schaefer et al., 2022), provide a possible explanation for some of the conflicting results in the literature (Schaefer et al., 2022). The stronger results in experiment 3 also shows that longer recording sessions (45 vs 5 min) increase the robustness of these patterns.

Although a clear pattern of pupil size fluctuations over the course of the breathing cycle emerges across averaged trials, this effect is not systematically evident at the level of individual breathing cycles. The cycle-by-cycle effects may be too subtle and thus masked by other dynamic factors that continuously influence momentary pupil size to a larger extent, such as cognitive, emotional, attentional, and arousal states (Mathôt, 2018).

Thus, determining the underlying mechanism and whether there is a cycle-by-cycle effect might require the use of animal models. These models would allow for more direct measurements and stricter control over a wider range of variables compared to human studies. However, the present study might offer some indirect insights. Breathing at rest (eupnea), as in our study, involves active inspiration, commonly followed by post-inspiration and passive expiration (Krohn et al., 2023; Negro et al., 2018). In this context, the most prevalent manner in which respiration can affect pupil size is through excitatory inputs from the preBötzinger complex to the locus coeruleus (LC), which are phase-locked to inspiration, leading to a respiration-regulated release of norepinephrine in the LC (Negro et al., 2018; Yackle et al., 2017). The LC projects to the intermedio-lateral column, which in turn projects to the superior cervical ganglion which innervates the iris dilator muscle (Mathôt, 2018; Szabadi, 2018).

Critically, as we observed the same pattern at rest and during a visual task, it is unlikely that the pupil dilation is caused by inhibition of the Edinger-Westphal nucleus parasympathetic constriction pathway. This inhibitory pathway is believed to be the primary pathway for dilation in response to arousal and mental effort (Mathôt, 2018; Szabadi, 2018). Therefore, we believe that the RPPE is not caused by cognitive, emotional, attentional, and arousal states per se. This is even more evident when comparing the RPPE to the spontaneous rhythmic constriction and dilation of the pupils known as hippus (Mathôt, 2018). The hippus is only present during rest and disappears during visual and cognitive tasks and increased arousal and so does its respiratory coupling (Kluger, Gross, & Keitel, 2023; Mathôt, 2018).

Importantly, nasal and oral breathing showed similar RPPE. This might indicate that the RPPE is shaped from the initial brainstem oscillators and not from the later olfactory bulb oscillator that is only activated when olfactory receptor neurons code nasal airflow and project this to the olfactory bulb. It should be noted that the olfactory bulb still might affect other aspects of the pupil function that have not been studied in this context, such as the complexity of the pupil dynamics (e.g., entropy and chaotic state).

Interestingly, we observed pupil dilation occurring throughout both inhalation and exhalation (see Figure 3). Therefore, even if the inspiratory signal from the preBötzinger complex is responsible for a drive towards pupil dilation, this drive does not abruptly end at exhalation onset. It would be of interest to investigate whether this pattern changes in scenarios in which exhalation is active rather than passive, such as during controlled breathing or exercise.

**Figure 3.**
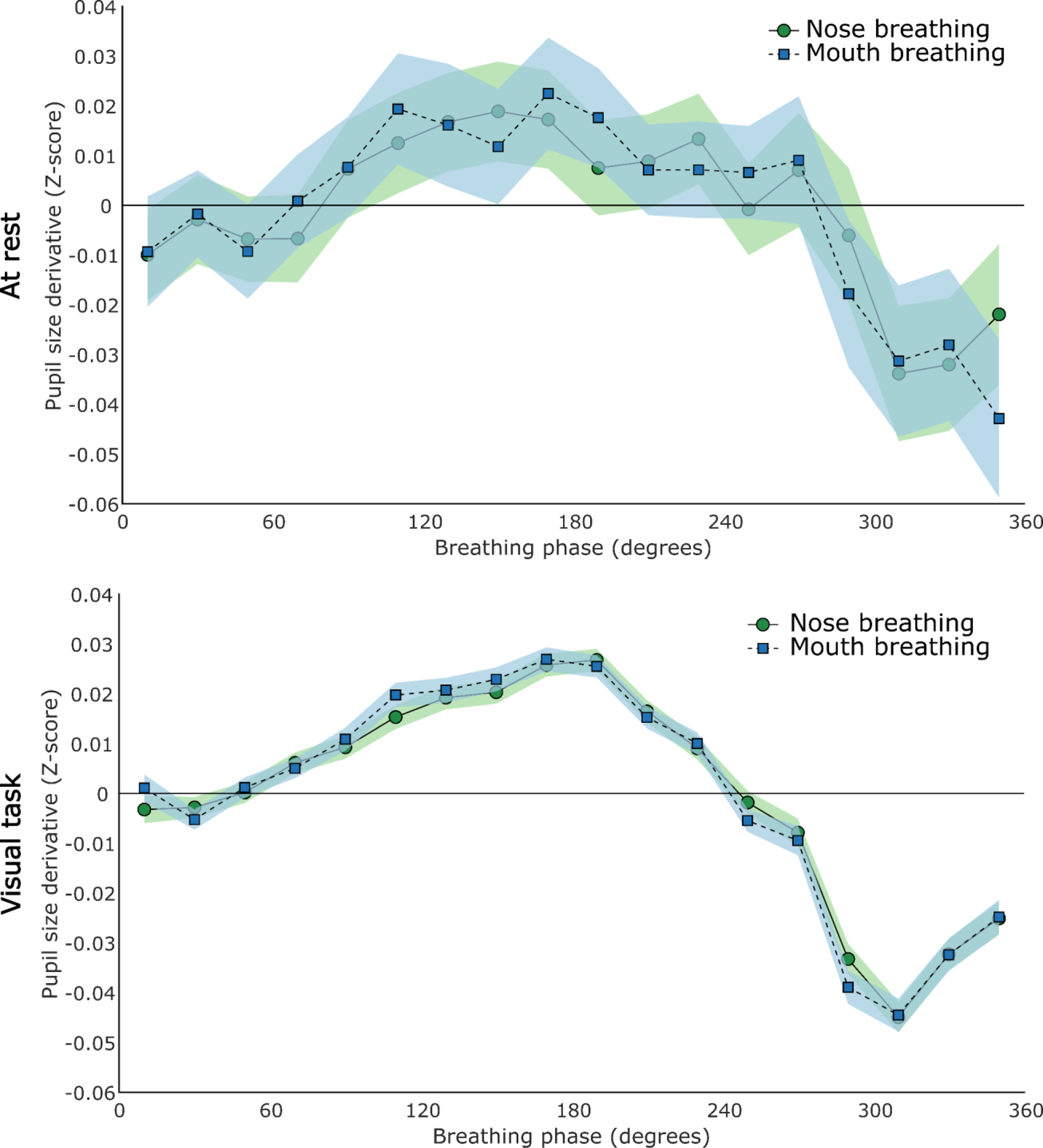
Average change in pupil size over the course of the breathing cycle. The line plot depicts the average change in normalized pupil size for each phase bin. The x-axis shows the breathing phase in degrees, with 0 indicating inhalation onset, and 180 indicating exhalation onset. The y-axis shows the change in normalized pupil size (derivative) in Z-scores. All values above the zero line indicate dilating pupils, and all values below the zero line indicate constricting pupils. Change in pupil size during nose breathing is depicted by the green circles connected by a solid line. Change in pupil size during mouth breathing is depicted by the blue squares connected by a dashed line. The shaded areas represent the 95% confidence intervals. The upper row depicts the results from the combined data recorded at rest, and the lower row the results from the Visual task data.

Future behavioral experiments should explore how these respiration-entrained fluctuations in pupil size impact visual perception. Past studies suggest a link between pupil size and task performance: smaller pupils benefit visual acuity tasks, while larger pupils aid in detecting faint stimuli (Mathôt & Ivanov, 2019). Our findings hint at the possibility that visual perception itself might cycle between optimizing for discrimination during inhalation and detection during exhalation within a single breath.

In conclusion, we have shown that, contrary to previous beliefs, pupil size is smallest around inhalation onset and largest during exhalation. This effect is robust and evident both during rest and active visual perception.

## Acknowledgements

We would like to thank Sylvia Edwards and Erik Gustavsson for their help with the data collection.

## Data and code availability

Preprocessed data and code will be made available upon journal publication. Raw data will be shared upon reasonable request.

