## Supplementary materials for "The Respiratory-Pupillary Phase Effect: Pupils size is smallest around inhalation onset and largest during exhalation"

| | At rest | | | | During visual task | | | |
|---|---|---|---|---|---|---|---|---|
| **Phase** | **Nose breathing** | | **Mouth breathing** | | **Nose breathing** | | **Mouth breathing** | |
| 0-20° | -0.048 | *p* < .001 | -0.072 | *p* < .001 | -0.064 | *p* < .001 | -0.060 | *p* < .001 |
| 20-40° | -0.060 | *p* < .001 | -0.067 | *p* < .001 | -0.069 | *p* < .001 | -0.066 | *p* < .001 |
| 40-60° | -0.066 | *p* < .001 | -0.065 | *p* < .001 | -0.070 | *p* < .001 | -0.066 | *p* < .001 |
| 60-80° | -0.069 | *p* < .001 | -0.066 | *p* < .001 | -0.062 | *p* < .001 | -0.057 | *p* < .001 |
| 80-100° | -0.045 | *p* < .001 | -0.045 | *p* < .001 | -0.054 | *p* < .001 | -0.047 | *p* < .001 |
| 100-120° | -0.014 | *p* < .001 | -0.015 | *p* < .001 | -0.037 | *p* < .001 | -0.029 | *p* < .001 |
| 120-140° | -0.012 | *p* < .001 | $1.73^{-4}$ | *p* = .955 | -0.028 | *p* < .001 | -0.019 | *p* < .001 |
| 140-160° | -0.003 | *p* = .464 | -0.006 | *p* = .068 | -0.019 | *p* < .001 | -0.011 | *p* < .001 |
| 160-180° | 0.012 | *p* < .001 | 0.014 | *p* < .001 | -0.004 | *p* < .001 | 0.013 | *p* < .001 |
| 180-200° | 0.010 | *p* = .005 | 0.016 | *p* < .001 | 0.053 | *p* < .001 | 0.060 | *p* < .001 |
| 200-220° | 0.020 | *p* < .001 | 0.028 | *p* < .001 | 0.079 | *p* < .001 | 0.081 | *p* < .001 |
| 220-240° | 0.035 | *p* < .001 | 0.037 | *p* < .001 | 0.090 | *p* < .001 | 0.089 | *p* < .001 |
| 240-260° | 0.023 | *p* < .001 | 0.048 | *p* < .001 | 0.089 | *p* < .001 | 0.082 | *p* < .001 |
| 260-280° | 0.045 | *p* < .001 | 0.047 | *p* < .001 | 0.079 | *p* < .001 | 0.082 | *p* < .001 |
| 280-300° | 0.062 | *p* < .001 | 0.045 | *p* < .001 | 0.050 | *p* < .001 | 0.047 | *p* < .001 |
| 300-320° | 0.039 | *p* < .001 | 0.037 | *p* < .001 | 0.005 | *p* < .001 | -0.010 | *p* < .001 |
| 320-340° | -0.011 | *p* = .001 | -0.016 | *p* < .001 | -0.036 | *p* < .001 | -0.045 | *p* < .001 |
| 340-360° | -0.041 | *p* < .001 | -0.035 | *p* < .001 | -0.060 | *p* < .001 | -0.067 | *p* < .001 |

Table S1. Average normalized pupil size (z-score) and *p*-values for each phase bin as calculated from the permutation testing results. The largest and smallest pupil size has been highlighted in gold and gray respectively, for each breathing route.

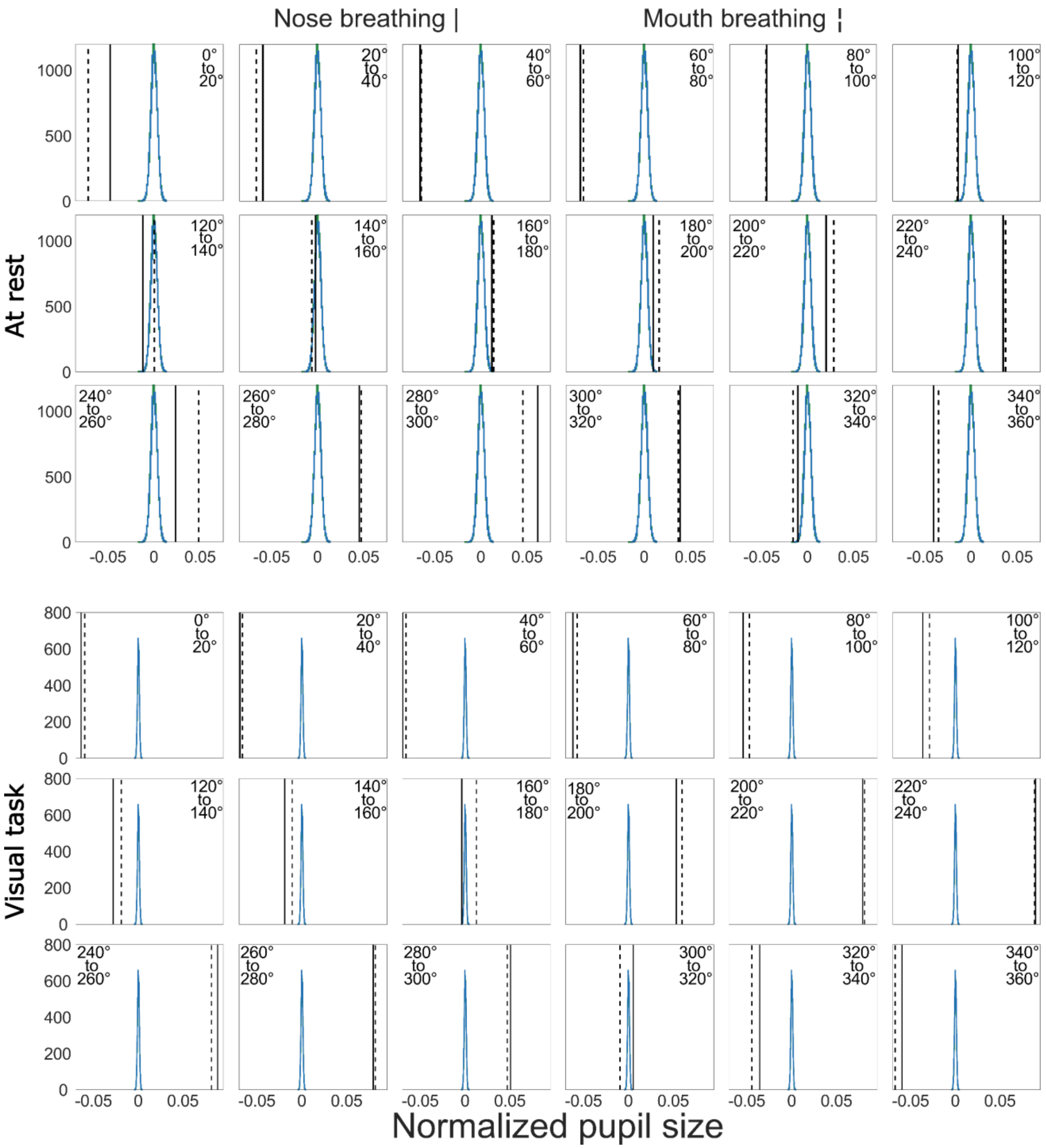


Figure S1 – Comparison of average pupil size during each of the 18 breathing bins to the distribution of means acquired by the permutation testing. The x-axis represents the average normalized pupil size in Z-scores, and the y-axis represents the number of occurrences of each pupil size acquired from the 10,000 permutations. The solid line represents the mean normalized pupil size for nose breathing, and the dashed line represents the same for mouth breathing. The green histogram shows the distribution of the permutation testing results for nose breathing, and the blue histogram shows the same for mouth breathing. Due to their high similarity, the two histograms overlap and are difficult to distinguish from each other. The upper row depicts the data from the combined recordings at rest (Experiment 1 & 2), and the lower row depicts the data from the recording during the visual task (Experiment 3).

Experiment 1

*Relative pupil size*

The results from the 2x2 repeated measures ANOVA showed a significant effect of breathing phase ($F(17) = 5.19$, $p < .001$, $\eta^2 = 0.079$), but no significant effect of breathing route ($F(1) = 0.45$, $p = .506$, $\eta^2 < 0.001$), nor a significant interaction effect ($F(17) = 0.63$, $p = .870$, $\eta^2 = 0.006$, see Figures S2 for graphical depictions).

The permutation testing revealed that 16/18 and 15/18 phase bins were significantly different from the randomly acquired distribution for nose and mouth breathing respectively, with 320° – 100/120° showing significantly smaller pupils, and 160/200° - 320° showing significantly larger pupils (see Table S2 and Figure S3 for the results of the permutation testing).

*Absolute pupil size*

The results from the 2x2 repeated measures ANOVA showed a significant effect of time ($F(17) = 3.97$, $p < .001$, $\eta^2 = 0.048$), but no significant effect of breathing route ($F(1) = 0.01$, $p = .939$, $\eta^2 < 0.001$), nor a significant interaction effect ($F(17) = 0.44$, $p = .976$, $\eta^2 = 0.003$, see Figure S4 for a graphical depiction of the results).

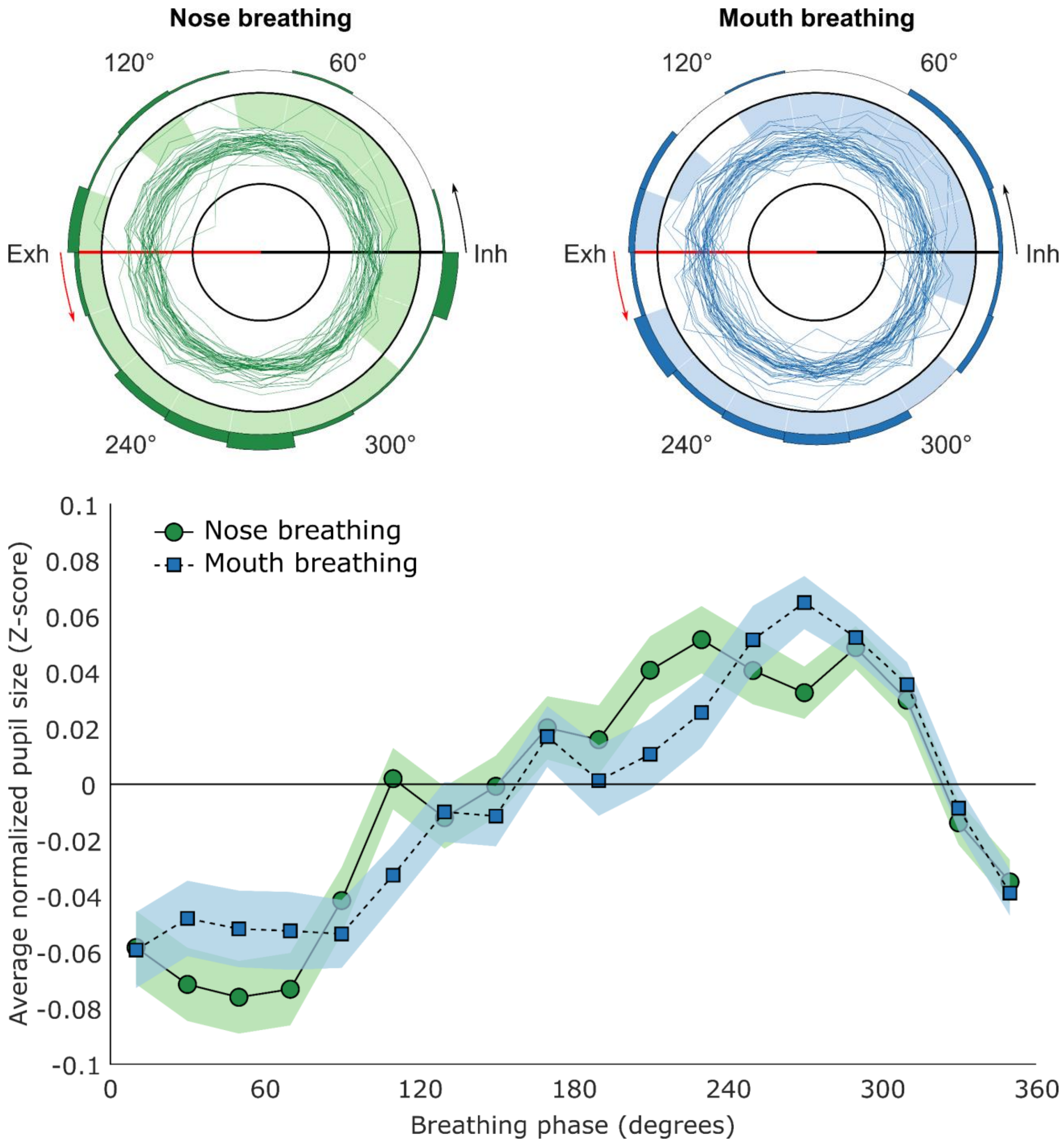


Figure S2 – The upper row shows the polar plots of participants' pupil size over the course of the breathing cycle (see Figure 1A legend for a more detailed description). The lower row depicts the average normalized pupil size for each phase bin. Zero degrees on the x-axis corresponds to inhalation onset, and 180° corresponds to exhalation onset (see Figure 1B legend for a more detailed description). Based on the data from Experiment 1.

The results from the one-way repeated measures ANOVAs showed a significant effect of breathing phase for nose breathing ($F(17) = 3.82$, $p < .001$, $\eta^2 = 0.087$), and for mouth breathing ($F(17) = 3.10$, $p < .001$, $\eta^2 = 0.067$).

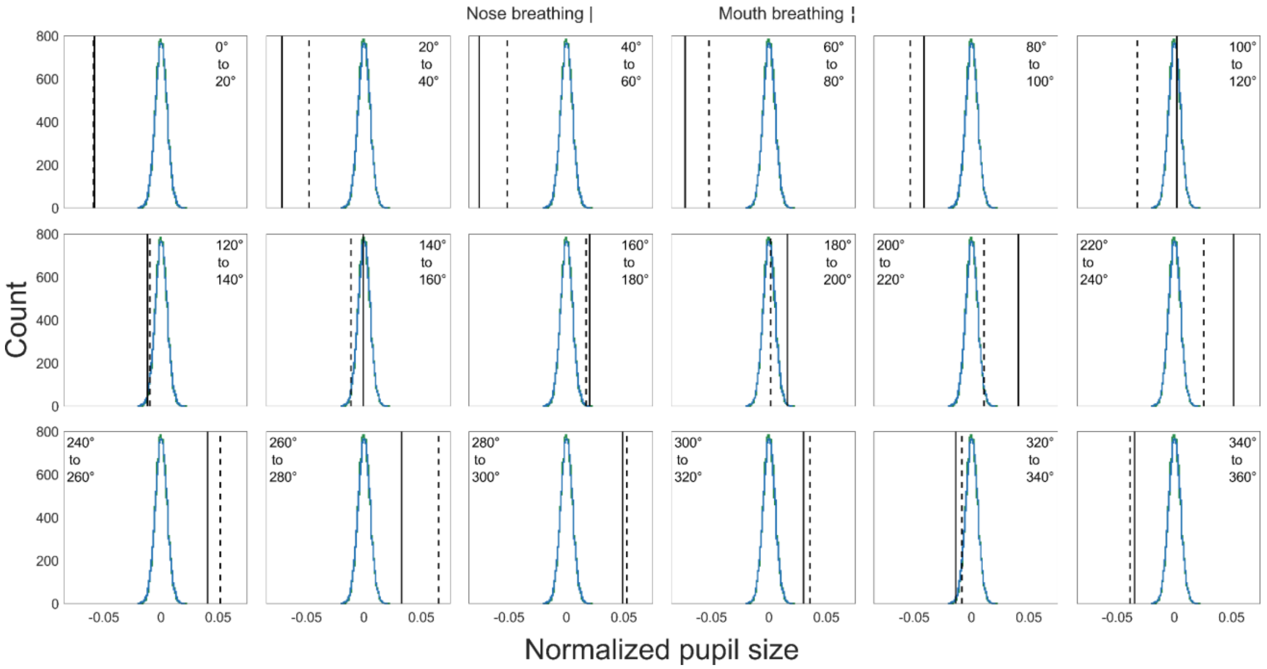


Figure S3 – Comparison of average pupil size during each of the 18 breathing bins to the distribution of means acquired by the permutation testing. The x-axis represents the average normalized pupil size in Z-scores, and the y-axis represents the number of occurrences of each pupil size acquired from the 10,000 permutations. The solid line represents the mean normalized pupil size for nose breathing, and the dashed line represents the same for mouth breathing. The green histogram shows the distribution of the permutation testing results for nose breathing, and the blue histogram shows the same for mouth breathing. Due to their high similarity, the two histograms overlap and are difficult to distinguish from each other. Based on the data from Experiment 1.

| **Phase** | **Nose breathing** | | **Mouth breathing** | |
|---|---|---|---|---|
| 0-20° | -0.058 | $p < .001$ | -0.059 | $p < .001$ |
| 20-40° | -0.072 | $p < .001$ | -0.048 | $p < .001$ |
| 40-60° | -0.076 | $p < .001$ | -0.052 | $p < .001$ |
| 60-80° | -0.073 | $p < .001$ | -0.052 | $p < .001$ |
| 80-100° | -0.042 | $p < .001$ | -0.054 | $p < .001$ |
| 100-120° | 0.002 | $p = .069$ | -0.033 | $p < .001$ |
| 120-140° | -0.012 | $p = .017$ | -0.010 | $p = .060$ |
| 140-160° | $-7.44^{-4}$ | $p = .888$ | -0.011 | $p = .029$ |
| 160-180° | 0.020 | $p < .001$ | 0.017 | $p < .001$ |
| 180-200° | 0.016 | $p = .002$ | 0.001 | $p = .810$ |
| 200-220° | 0.041 | $p < .001$ | 0.011 | $p = .042$ |

| | | | | |
|---|---|---|---|---|
| 220-240° | 0.052 | $p < .001$ | 0.026 | $p < .001$ |
| 240-260° | 0.041 | $p < .001$ | 0.052 | $p < .001$ |
| 260-280° | 0.033 | $p < .001$ | 0.065 | $p < .001$ |
| 280-300° | 0.049 | $p < .001$ | 0.052 | $p < .001$ |
| 300-320° | 0.030 | $p < .001$ | 0.036 | $p < .001$ |
| 320-340° | -0.014 | $p = .006$ | -0.009 | $p = .107$ |
| 340-360° | -0.035 | $p < .001$ | -0.040 | $p < .001$ |

Table S2 - Average normalized pupil size and *p*-values for each phase bin as calculated from the permutation testing results, based on the data from Experiment 1.

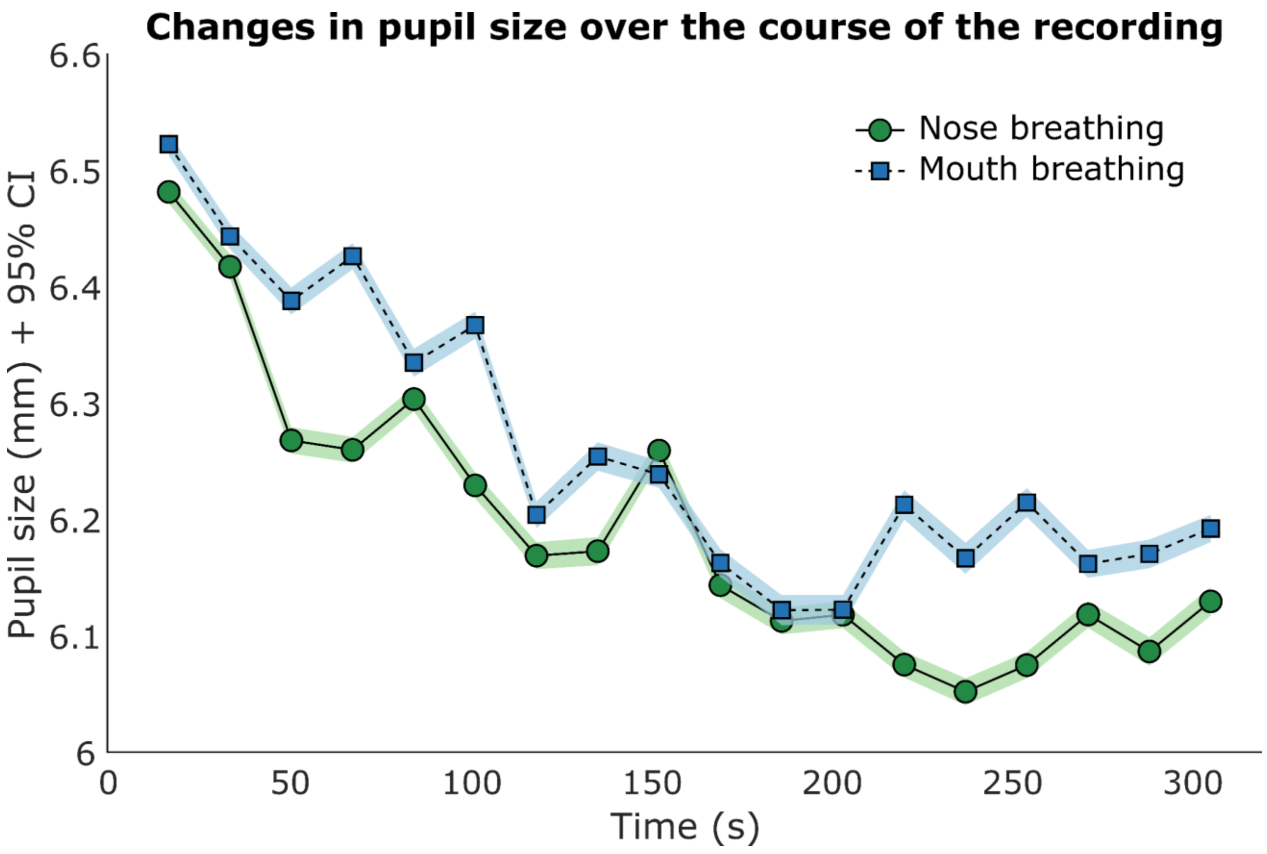


Figure S4 - Average pupil size over the course of the recording. The line plot depicts the average absolute pupil size for each phase bin. The x-axis shows the recording time in seconds. The y-axis shows pupil size in mm. Pupil size during nose breathing is depicted by the green circles connected by a solid line, and pupil size during mouth breathing is depicted by the blue squares connected by a dashed line. The error bars represent the 95% confidence interval. Based on the data from Experiment 1.

Experiment 2

*Relative pupil size*

The results from the 2x2 repeated measures ANOVA showed a significant effect of breathing phase ($F(17) = 5.18$, $p < .001$, $\eta^2 = 0.064$), but no significant effect of breathing route ($F(1) = 3.30$, $p = .076$, $\eta^2 = 0.001$), nor a significant interaction effect ($F(17) = 0.87$, $p = .611$, $\eta^2 = 0.008$, see Figures S5 for graphical depictions).

The permutation testing revealed that 12/18 and 15/18 phase bins were significantly different from the randomly acquired distribution for nose and mouth breathing respectively, with 320/340° – 100/140° showing significantly smaller pupils, and 160/260° - 320° showing significantly larger pupils (see Table S3 and Figure S6 for the results of the permutation testing).

*Absolute pupil size*

The results from the 2x2 repeated measures ANOVA showed a significant effect of time ($F(17) = 4.64$, $p < .001$, $\eta^2 = 0.038$), but no significant effect of breathing route ($F(1) = 1.18$, $p = .284$, $\eta^2 = 0.012$), nor a significant interaction effect ($F(17) = 0.76$, $p = .740$, $\eta^2 = 0.004$, see Figure S7 for a graphical depiction of the results).

When combining the data from Experiment 1 & 2, the results did not change with the 2x2 repeated measures ANOVA showing a significant effect of time ($F(17) = 8.11$, $p < .001$, $\eta^2 = 0.040$), but no significant effect of breathing route ($F(1) = 0.77$, $p = .383$, $\eta^2 = 0.004$), nor a significant interaction effect ($F(17) = 0.84$, $p = .651$, $\eta^2 = 0.003$).

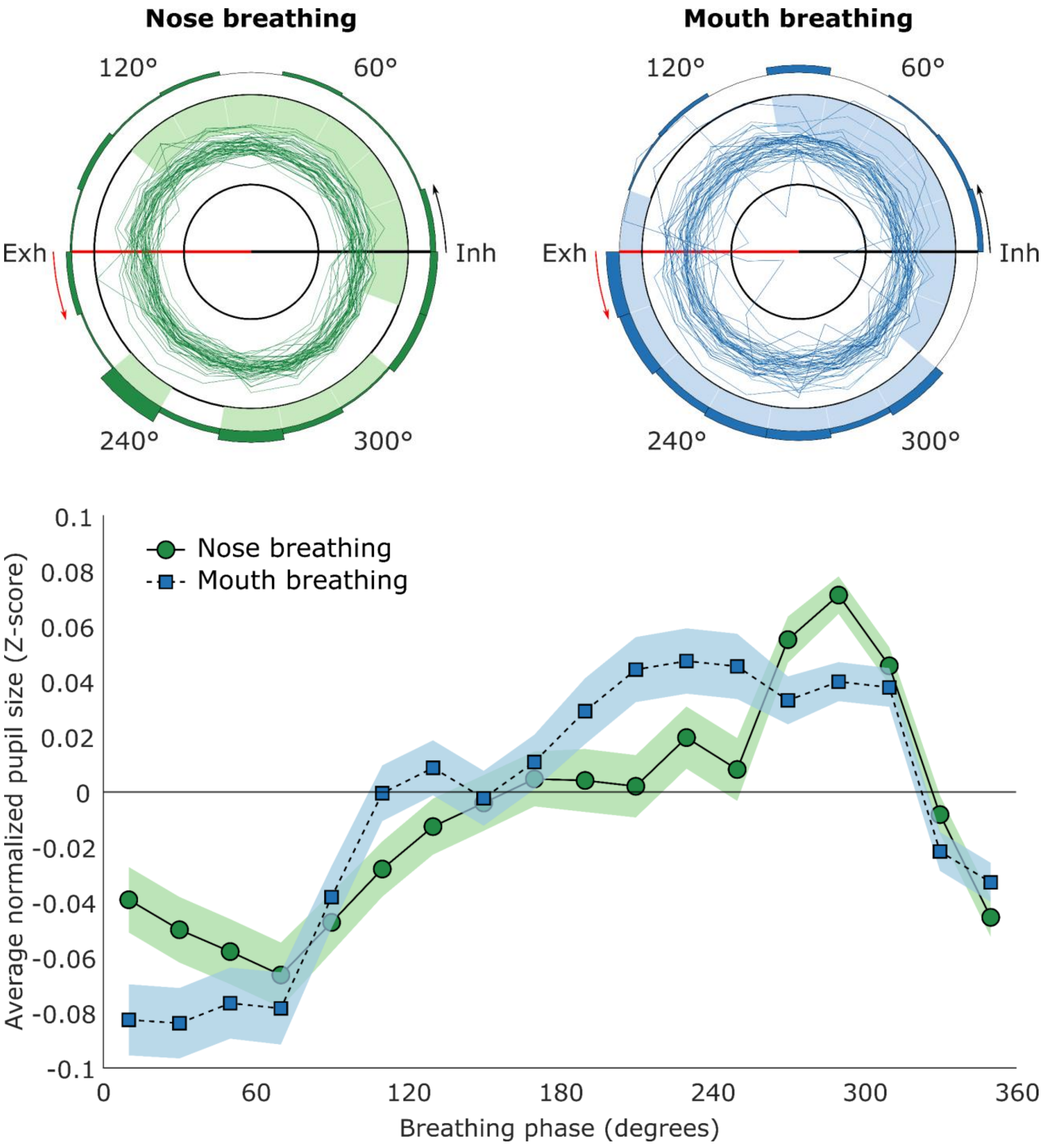


Figure S5 - The upper row shows the polar plots of participants' pupil size over the course of the breathing cycle (see Figure 1A legend for a more detailed description). The lower row depicts the average normalized pupil size for each phase bin. Zero degrees on the x-axis corresponds to inhalation onset, and 180° corresponds to exhalation onset (see Figure 1B legend for a more detailed description). Based on the data from Experiment 2.

The results from the one-way repeated measures ANOVAs showed a significant effect of breathing phase for nose breathing ($F(17) = 5.33$, $p < .001$, $\eta^2 = 0.102$), and for mouth breathing ($F(17) = 3.18$, $p < .001$, $\eta^2 = 0.065$).

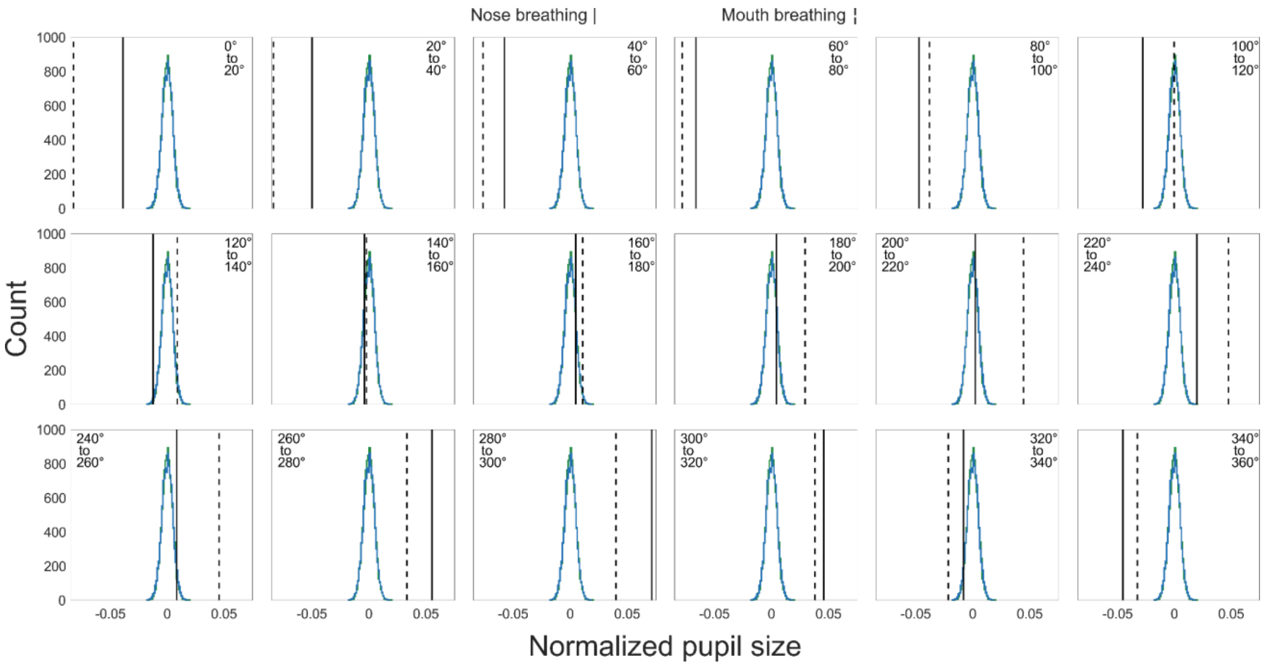


Figure S6 - Comparison of average pupil size during each of the 18 breathing bins to the distribution of means acquired by the permutation testing. The x-axis represents the average normalized pupil size in Z-scores, and the y-axis represents the number of occurrences of each pupil size acquired from the 10,000 permutations. The solid line represents the mean normalized pupil size for nose breathing, and the dashed line represents the same for mouth breathing. The green histogram shows the distribution of the permutation testing results for nose breathing, and the blue histogram shows the same for mouth breathing. Due to their high similarity, the two histograms overlap and are difficult to distinguish from each other. Based on the data from Experiment 2.

| Phase | Nose breathing | | Mouth breathing | |
|---|---|---|---|---|
| 0-20° | -0.039 | $p < .001$ | -0.083 | $p < .001$ |
| 20-40° | -0.050 | $p < .001$ | -0.084 | $p < .001$ |
| 40-60° | -0.058 | $p < .001$ | -0.077 | $p < .001$ |
| 60-80° | -0.066 | $p < .001$ | -0.078 | $p < .001$ |
| 80-100° | -0.047 | $p < .001$ | -0.038 | $p < .001$ |
| 100-120° | -0.028 | $p < .001$ | $-4.64^{-4}$ | $p = .919$ |
| 120-140° | -0.013 | $p = .005$ | 0.009 | $p = .067$ |
| 140-160° | -0.004 | $p = .392$ | -0.002 | $p = .642$ |
| 160-180° | 0.005 | $p = .306$ | 0.011 | $p = .022$ |

| | | | | |
|---|---|---|---|---|
| 180-200° | 0.004 | *p* = .367 | 0.029 | *p* < .001 |
| 200-220° | 0.002 | *p* = .648 | 0.044 | *p* < .001 |
| 220-240° | 0.020 | *p* < .001 | 0.047 | *p* < .001 |
| 240-260° | 0.008 | *p* = .069 | 0.046 | *p* < .001 |
| 260-280° | 0.055 | *p* < .001 | 0.033 | *p* < .001 |
| 280-300° | 0.071 | *p* < .001 | 0.040 | *p* < .001 |
| 300-320° | 0.046 | *p* < .001 | 0.038 | *p* < .001 |
| 320-340° | -0.008 | *p* = .078 | -0.022 | *p* < .001 |
| 340-360° | -0.045 | *p* < .001 | -0.033 | *p* < .001 |

Table S3 - Average normalized pupil size and *p*-values for each phase bin as calculated from the permutation testing results, based on the data from Experiment 2.

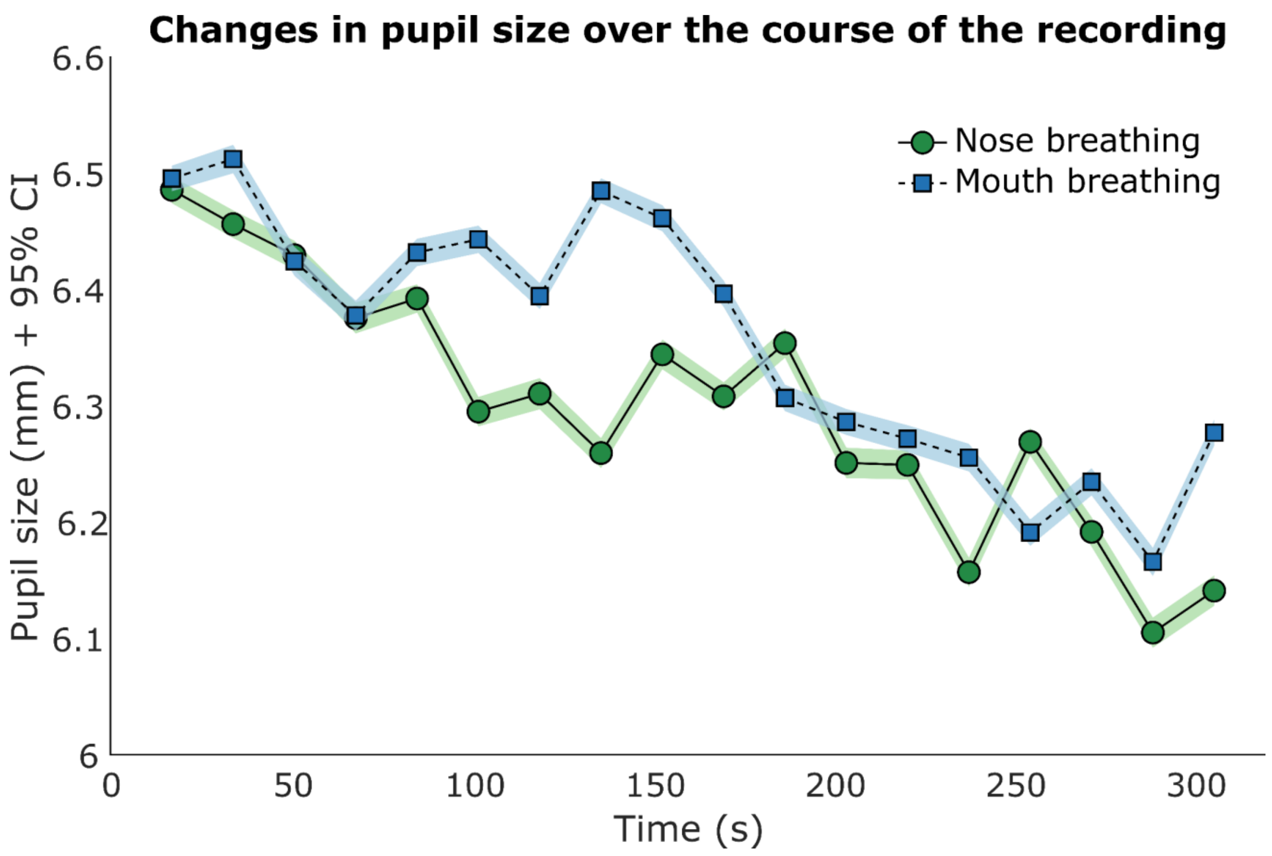


Figure S7 - Average pupil size over the course of the recording. The line plot depicts the average absolute pupil size for each phase bin. The x-axis shows the recording time in seconds. The y-axis shows pupil size in mm. Pupil size during nose breathing is depicted by the green circles connected by a solid line, and pupil size during mouth breathing is depicted by the blue squares connected by a dashed line. The error bars represent the 95% confidence interval. Based on the data from Experiment 2.
